# Nuclei multiplexing with barcoded antibodies for single-nucleus genomics

**DOI:** 10.1101/476036

**Authors:** Jellert T. Gaublomme, Bo Li, Cristin McCabe, Abigail Knecht, Eugene Drokhlyansky, Nicholas Van Wittenberghe, Julia Waldman, Danielle Dionne, Lan Nguyen, Phil De Jager, Bertrand Yeung, Xinfang Zhao, Naomi Habib, Orit Rozenblatt-Rosen, Aviv Regev

**Affiliations:** Klarman Cell Observatory, Broad Institute of Harvard and MIT, Cambridge, MA, USA.; Center for Translational & Computational Neuroimmunology, Columbia University Medical Center, New York, NY, USA.; BioLegend Inc., San Diego, CA, USA; Edmond and Lily Safra Center for Brain Sciences, Hebrew University of Jerusalem, Jerusalem, Israel; Howard Hughes Medical Institute, Koch Institute of Integrative Cancer Research, Department of Biology, Massachusetts Institute of Technology, Cambridge, MA, USA

## Abstract

Single-nucleus RNA-Seq (snRNA-seq) enables the interrogation of cellular states in complex tissues that are challenging to dissociate, including frozen clinical samples. This opens the way, in principle, to large studies, such as those required for human genetics, clinical trials, or precise cell atlases of large organs. However, such applications are currently limited by batch effects, sequential processing, and costs. To address these challenges, we present an approach for multiplexing snRNA-seq, using sample-barcoded antibodies against the nuclear pore complex to uniquely label nuclei from distinct samples. Comparing human brain cortex samples profiled in multiplex with or without hashing antibodies, we demonstrate that nucleus hashing does not significantly alter the recovered transcriptome profiles. We further developed demuxEM, a novel computational tool that robustly detects inter-sample nucleus multiplets and assigns singlets to their samples of origin by antibody barcodes, and validated its accuracy using gender-specific gene expression, species-mixing and natural genetic variation. Nucleus hashing significantly reduces cost per nucleus, recovering up to about 5 times as many single nuclei per microfluidc channel. Our approach provides a robust technique for diverse studies including tissue atlases of isogenic model organisms or from a single larger human organ, multiple biopsies or longitudinal samples of one donor, and large-scale perturbation screens.

## Introduction

Single-nucleus RNA-seq (snRNA-Seq) has become an instrumental method for interrogating cell types, states, and function in complex tissues that cannot easily be dissociated (*1–3*). This includes tissues rich in cell types such as neurons, adipocytes and skeletal muscle cells, archived frozen clinical materials, and tissues that must be frozen to register into specific coordinates. Moreover, the ability to handle minute frozen specimens (*4*) has made snRNA-seq a compelling option for large scale studies from tissue atlases (*5, 6*), to longitudinal clinical trials and human genetics. However, to maximize the success of such studies there is a crucial need to minimize batch effects, reduce costs, and streamline the preparation of large numbers of samples.

For single *cell* analysis, these goals have recently been elegantly achieved by multiplexing samples prior to processing, which are barcoded either through natural genetic variation (*7*), chemical labeling (*8, 9*) or DNA-tagged antibodies (“cell hashing”) (*10*). These methods have improved technical inter-sample variability by early pooling, lower the cost per sample by overloading cells per microfluidic run — due to an increased ability to detect and discard co-encapsulated “cell multiplets” sharing the same bead barcode — and reduce the number of parallel processing steps in large studies.

Here, we follow on these studies by developing a sample multiplexing method for nuclei (“nucleus hashing”), using DNA-barcoded antibodies targeting the nuclear pore complex. Unlike methods leveraging natural genetic variation (*7*), barcoded antibodies allow pooling of isogenic samples, such as from isogenic mouse models, multiple specimens from the same human donor, or tissues sampled and preserved from a given donor over time.

## Results

We isolated nuclei from fresh-frozen murine or human cortical tissues, stained them with antibodies carrying a sample-specific DNA barcode, and pooled samples prior to droplet encapsulation for single-nucleus RNA-Seq (snRNA-Seq) (**Figure 1a**). The DNA barcodes contain a polyA tail, thus acting as artificial transcripts that register the same bead barcode as nuclear transcripts, coupling the transcription profile to the sample of origin.

**Figure 1.**
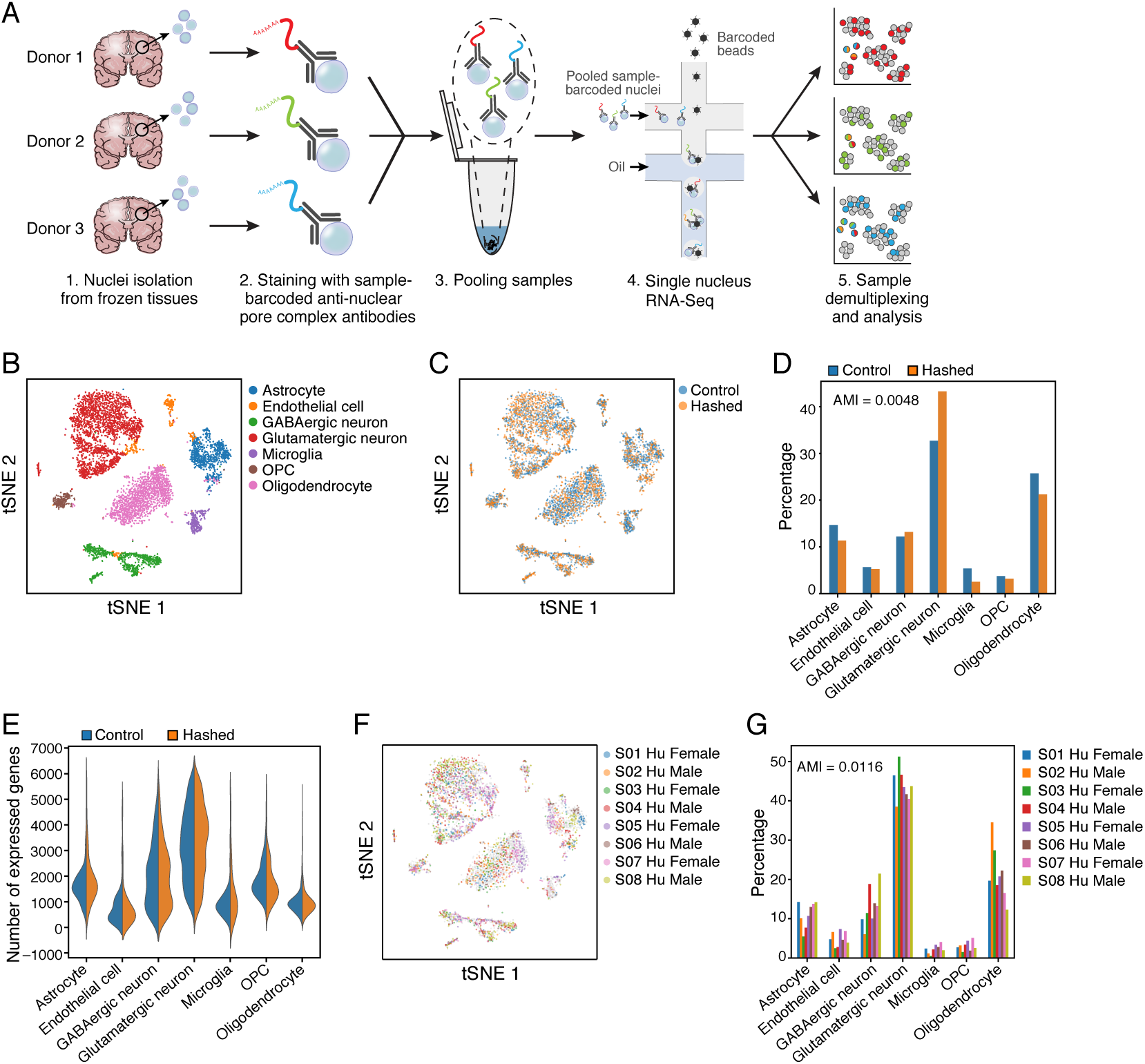
Nuclei multiplexing using DNA-barcoded antibodies targeting the nuclear pore complex. **a.** Experimental workflow. Nuclei are isolated from frozen tissues and stained with DNA-barcoded antibodies targeting the nuclear pore complex (MAb414, Biolegend). The DNA barcode encodes a unique sequence representing each tissue sample, enabling sequence-based identification of each nucleus after pooling and profiling the different samples. **b-e.** Multiplexed and non-multiplexed samples of human cortex from 8 postmortem donors yield comparable results. **b.** *t*-stochastic neighborhood embedding (tSNE) of single nucleus profiles (dots) colored by either cell type (**b**) or by type of protocol (**c**). Non-hashed control sample (blue) and hashed sample (orange) show similar patterns. **d.** Cell type frequencies observed for hashed (orange) and non-hashed control (blue) samples. The adjusted mutual information (AMI) is shown in the top left. **e.** Distributions of the number of expressed genes (*y* axis, left) in each cell type (*x* axis) in **b**, for nuclei from hashed (orange) and non-hashed control (blue) samples. **f-g.** Hashed single nuclei from all donors are similarly represented across cell type clusters. **f.** tSNE as in **b** colored by donor. **g.** Observed frequencies (*y* axis) of each cell type (*x* axis) per donor (color). The adjusted mutual information (AMI) is shown in the top left.

The additional antibody labeling step in our protocol did not alter the quality of transcriptional profiling compared to non-hashed snRNA-seq, in a side-by-side comparison of a hashed (antibody labeled) *vs*. non-hashed pool of cortex nuclei derived from eight human donors (**Supplementary Table 1**). We combined the expression profiles from both hashed and non-hashed datasets, followed by clustering and *post-hoc* annotation with legacy cell type-specific signatures (**Figure 1b**), recovering all cell types previously reported for such samples (*1*) (**Methods**). Both hashed and non-hashed nuclei were similarly represented across the recovered cell types (**Figure 1c**), with an adjusted mutual information score of 0.0048 between cell types and experimental conditions (**Figure 1d**, **Methods**), with only slight differences, such as a weak enrichment of glutamatergic neurons in the hashed sample, and similar cell type-specific numbers of recovered genes (**Figure 1e**). Each cell type had nuclei from all 8 donors (**Figure 1f**) with only slightly differing frequencies (**Figure 1g**), as expected for a diverse donor cohort (*1*) (**Supplementary Table 1**). Notably, modifying the staining and washing buffers for nucleus hashing (**Methods**) compared to those used in cell hashing (*10*) improved the transcriptional similarity with the non-hashed control (**Supplementary Figure 1a**), and achieved a similar number of genes expressed per nucleus as the non-hashed control (**Supplementary Figure 1b**), whereas a PBS based buffer (used in cell hashing (*10*)) generally had poorer performance (**Supplementary Figure 1c**). We thus performed all experiments with these novel staining and washing buffers, except those with mouse samples. Collectively, these findings indicate that hashing preserves library quality and cell type distributions.

To probabilistically assign each nucleus to its sample barcode, we developed DemuxEM, an Expectation-Maximization-based tool (**Figure 2a**). For each nucleus, DemuxEM takes as input a vector of hashtag Unique Molecular Identifiers (UMIs) from that nucleus (**Figure 2a**, left). The input vector is a mixture of signal hashtag UMIs, which reflect the nucleus’ sample of origin, and background hashtag UMIs, which likely reflect ambient sample barcodes. Hashtag UMIs from the background have different probabilities of matching each of the sample barcodes. DemuxEM estimates this background distribution of sample barcodes based on hashtag UMIs in empty droplets, which are likely to only contain background hashtag UMIs. With this background distribution as a reference, DemuxEM uses an Expectation-Maximization (EM) algorithm to estimate the fraction of hashtag UMIs from the background in the given droplet and then infer the signal hashtag UMIs by deducting the estimated background UMIs from the input vector. Once the signal has been identified, DemuxEM determines if this droplet encapsulated a single nucleus or a multiplet. For bead barcodes with low signal hashtag UMIs (*e.g.*, < 10 hashtag UMIs), DemuxEM cannot determine the origin of the nucleus and marks it as ‘unassigned’ (**Methods**).

**Figure 2.**
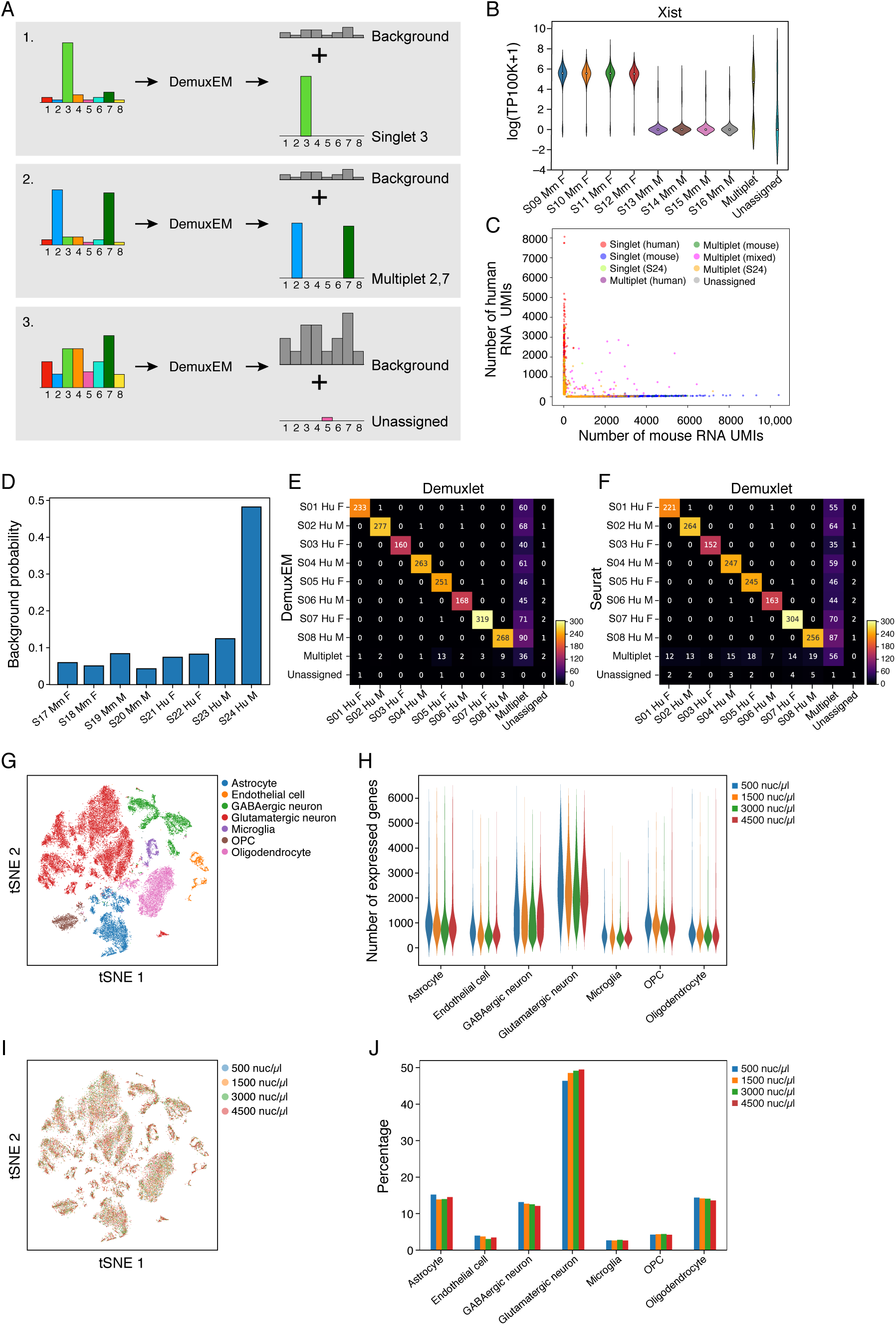
Accurate sample assignment by DemuxEM allows efficient overloading of hashed samples. **a.** Sample assignment by DemuxEM. DemuxEM takes as input for each nucleus a count vector of hashtag UMIs (left) and estimates them as a sum of a background hashtag UMI vector in that nucleus (right, grey histograms) and a signal sample assignment hashtag UMI vector (right, color histograms). Shown are schematic examples: singlet assignment (top), multiplet detection (middle), and unassigned (bottom). **b.** Validation of DemuxEM assignment by gender mixing in isogenic mice. Distribution of Xist expression (*y* axis, log(TP100K+1)) from 8 mouse-derived cortex samples (samples 1-4 female, samples 5-8 male) that were pooled and demultiplexed. There is 94.8% agreement between DemuxEM-assigned sample hashtag identities and Xist expression among DemuxEM-detected singlets. **c,d.** DemuxEM assignments in species mixing of human and mouse cortex nuclei. **c.** Species mixing plot. Each nucleus (dot) is plotted by the number of RNA UMIs aligned to pre-mRNA mouse mm10 (*x* axis) and human GRCh38 (*y* axis) references (**Methods**), and colored by its DemuxEM-predicted hashtag sample identities for singlet human (red), singlet mouse (blue) or different multiplets (intra-species: green (mouse) and purple (human); inter-species: fuchsia). S24 singlets (chartreuse) and multiplets (orange) are colored separately due to its large contribution to ambient hashtags. **d.** Distribution of ambient hashtags matching the sample DNA barcode (x-axis) in the pool of 8 samples. DemuxEM identified S24 as a disproportionate contributor to the hashtag background distribution. **e,f.** Validation of hashtag-based assignment of nuclei by natural genetic variation. Shown is the number of nuclei classified as sample singlet, multiplets or unassigned (rows, columns) by either natural genetic variation (columns) with Demuxlet (*7*), or based on hashtag UMIs (rows), with DemuxEM (**e**) or Seurat (*11*) (**f**). 98.1% of nuclei identified by Demuxlet as singlets from a given donor are similarly identified by DemuxEM, and hashtag-based classification recovers more singlets than by natural variation. **g-j.** Nucleus hashing allows over-loading to reduce experimental costs. **g.** tSNE of combined data of 8 hashed human cortex samples profiled by snRNA-Seq at loading concentrations of 500, 1,500, 3,000 or 4,500 nuclei/μl. Single nucleus profiles (dots) are colored by cell type. **h.** Comparable distributions of the number of expressed genes (*y* axis) in each cell type (*x* axis) in **g**, for nuclei from each loading density. **i.** tSNE of single nucleus profiles (dots) as in **g**, colored by loading concentration. **j.** Comparable frequencies (y axis) across cell types in **g** (x axis) observed for different loading concentrations.

To assess our confidence in calling the sample origin of hashed nuclei by their sample barcodes, we next applied DemuxEM to pooled nuclei of male and female isogenic mice or of human and mouse, such that the single nucleus transcriptomes provided an orthogonal measure of the sample of origin. First, we multiplexed nuclei isolated from two isogenic C57BL/6J mouse cortices, 4 technical replicates from each of a female and male mouse (**Methods**). For DemuxEM-identified singlets, there was a 94.8% agreement between DemuxEM-assigned sample hashtag identities and the expression level of Xist, a transcript predominantly expressed in females (**Figure 2b**). Next, we multiplexed 8 cortex samples, 4 from mouse and 4 from human (**Supplementary Table 1**), comparing DemuxEM assignment as human or mouse singlets to their position in a “species-mixing plot” based on their number of RNA UMIs mapping to the human or mouse transcriptome (**Methods**, **Figure 2c**). Overall, nuclei assigned by DemuxEM as human or mouse singlets (**Figure 2c**, red and blue, respectively) express predominantly human or mouse reads, respectively (**Figure 2c**, alignment along the Y and X axis). DemuxEM-predicted multiplets occur both on the species-specific axes for intra-species multiplets (**Figure 2c**, green (mouse) and purple (human)) and off-axes for inter-species multiplets (**Figure 2c**, fuchsia).

We further leveraged the hashtags to address the sources of ambient hashtags in a pool of samples. In general, nuclei dissociated from tissue samples may be at risk of having higher levels of ambient hashtags compared to single-cell hashing, because the cytoplasm is disrupted during lysis and nonspecific antibody binding to cytosolic content or tissue derived debris could contribute to the background. Inspection of sample-specific contribution to the hashtag background distribution showed that one of the human samples (S24, **Supplementary Table 1**) contributed disproportionally to the background (**Figure 2d**), suggesting that this sample might have been of lower quality. This donor sample (S24) indeed had the lowest RNA integrity number (RIN) and the highest post mortem interval (PMI) of all subjects in the study (**Supplementary Table 1**) The ability to identify which samples contribute to the background is an additional benefit of sample hashing, and can help determine quality parameters for sample inclusion.

Next, we validated our hashtag based demultiplexing with Demuxlet (*7*), an approach based on natural genetic variation. We observed excellent agreement between the two methods for the 8 human cortex samples (**Figure 2e**): on average, 98.1% of the nuclei identified by Demuxlet as single nuclei from a given donor are similarly identified by DemuxEM (**Figure 2e**). Moreover, demultiplexing based on the hashtag data enables the identification of more singlets per donor when using either DemuxEM or Seurat, a package that includes single-cell hashing analysis (*11*) (**Figure 2e,f**, **Supplementary Table 2**).

DemuxEM also offers a better estimation of the multiplet rate. The multiplet rate per 10X microfluidic channel when loading 7,000 cells is expected to be ~3.1% (*12*). When pooling 8 samples with equal proportions, there are 56 possible *inter*-sample doublet configurations and 8 possible intra-sample ones (the proportion of higher order multiplets is much lower), such that 87.5% (56/64) of the doublets are expected to contain nuclei from multiple samples, which can be identified by our hashing strategy. Since we loaded 7,000 nuclei, we expect a detectable multiplet rate of at least 2.7% (3.1 * 87.5%). DemuxEM, Seurat, and Demuxlet predicted multiplet rates of 2.8%, 6.5%, and 20.6%, respectively (**Supplementary Table 2**).

This ability to more accurately detect droplets that encapsulated multiple inter-sample nuclei allowed us to load a higher concentration of nuclei for a given undetectable multiplet rate, thereby significantly lowering the cost per nucleus. To assess how ‘over-loading’ a higher concentration of nuclei affects library quality and cell type distributions, we hashed and pooled another 8 human cortex samples (**Supplementary Table 1**) and loaded a 10X channel with 14 μl of either ~500 nuclei/μl, 1,500 nuclei/μl, 3,000 nuclei/μl or 4,500 nuclei/μl. When sequencing these libraries at similar depth *per nucleus*, we recovered similar numbers of expressed genes per nucleus for the different cell types (**Figure 2g,h**). Moreover, nuclei from each loading concentration had similar transcriptional states (**Figure 2i**) and maintained the same relative cell type frequencies (**Figure 2j**). As expected, the proportion of multiplets increases with increased loading density (**Supplementary Figure 2**). Notably, nucleus multiplets do not typically show higher numbers of RNA UMIs compared to singlets (**Supplementary Figure 2**), in contrast to cell-hashing (*10*). The lowest overall cost per nucleus (including nucleus-hashing antibodies, 10X library preparation and sequencing) was achieved for loading 14μl of 3,000 nuclei/μl, resulting in the detection of 13,578 single nuclei in a single 10X channel with an overall ~56% cost per nucleus reduction in our pricing structure, compared to the non-hashed loading density of 500 nuclei/μl (**Methods, Supplementary Table 3**), albeit with some increase in background signal. Notably, these cost savings can also be achieved by splitting an *individual* sample into multiple hashed samples, when a larger number of nuclei per sample is required, while still benefitting from the reduced cost and reduced batch effects.

## Discussion

Nucleus hashing is a principled method for multiplexing single nuclei. It reduces batch effects and costs and helps streamline large experimental studies. DemuxEM is a novel computational tool that enables accurate multiplet detection, nucleus identity assignment, and identification of the sources of ambient hashtag contamination. As nuclei, rather than cells, become the starting point of many additional methods – especially in epigenomics – it is likely that hashing can be extended to other single nucleus genomics assays. Together, nucleus hashing and DemuxEM allow us to reliably interrogate cell types, cellular states, and functional processes in complex and archived tissues at a much larger scale than previously possible.

## Supporting information

## Data availability

**All mouse data** will be available from the Gene Expression Omnibus and the Single Cell Portal: https://portals.broadinstitute.org/single_cell.

**Processed human expression** data will be available from the Gene Expression Omnibus and in the Single Cell Portal: https://portals.broadinstitute.org/single_cell.

**Raw human sequencing** data will be available in the Broad DUOS system: https://duos.broadinstitute.org/#/home.

**DemuxEM** will be released as part of the scCloud for single cell analysis, available at https://github.com/klarman-cell-observatory/KCO

**Whole Genome Sequencing** data for the human samples can be obtained through the AMP-AD Knowledge Portal that is supported by the National Institute of Aging (https://www.synapse.org/#!Synapse:syn2580853/wiki/409840).

## Acknowledgments

We thank Dr. David Bennett at RUSH University for the use of samples from the Religious Order Study (ROS) and the Memory and Aging Project (MAP). We thank Leslie Gaffney, Anna Hupalowska and Jennifer Rood for help with figure and paper preparation. This work was supported by the NIH BRAIN Initiative grant U19MH114821 and the Klarman Cell Observatory. The ROSMAP sample collection and data used in this manuscript were supported by U01 AG046152, RF1 AG057473, and RF1 AG015819.

## Author Contributions

J.T.G. and A.R. conceived the study and designed experiments. B.L., and A.R devised analyses and B.L. developed computational methods. B.L., J.T.G. and A.R analyzed the data. X.Z and B.Y validated and provided hashing antibodies. J.T.G., C.M., N.V.W, E.D, A.K and J.W. conducted the experiments. L.N, J.W, J.T.G and D.D. carried out Illumina library preparation. O.R.R. and A.R. supervised work. J.T.G, B.L. and A.R. wrote the paper with input from all the authors.

## Materials and Methods

**Human samples.** The study was conducted under IRB approval L91020181. We used frozen brain tissue from the dorsolateral prefrontal cortex (DLPFC) banked by two prospective studies of aging: the Religious Order Study (ROS) and the Memory and Aging Project (MAP), which recruit non-demented older individuals (age >65) (*13*). We selected samples for which Whole Genome Sequencing data was already available (*14*). We selected 10 males and 10 females (**Supplementary Table 1**).

### Mice

All mouse work was performed in accordance with the Institutional Animal Care and Use Committees (IACUC) and relevant guidelines at the Broad Institute and MIT, with protocol 0122-10-16. Adult female and male C57BL/6J mice, obtained from the Jackson Laboratory (Bar Harbor, ME), were housed under specific-pathogen-free (SPF) conditions at the Broad Institute, MIT animal facilities.

### Mouse tissue collection

Brains from C57BL/6J mice were obtained and split vertically along the sagittal midline. The cerebral cortices were separated and excess white matter was removed. Cortices were separated into microcentrifuge tubes and frozen on dry ice. Frozen tissue was stored at −80 °C.

### Nuclei isolation, antibody tagging, and snRNA-seq

A fully detailed, step-by-step protocol, is described in as the **Protocol** below. Briefly, we thawed and minced tissue, dounced it in lysis buffer, filtered the lysate, and resuspended it in staining buffer. A brief incubation with Fc receptor blocking solution is followed by incubation with the TotalSeq Hashtag antibodies and 3 washes in ST-SB. Next, nuclei were counted and their concentration normalized to the desired loading concentration and pooled right before running the 10X Genomics single-cell 3’ v2 assay (with minor adjustments listed in the detailed protocol), followed by library preparation and Illumina sequencing.

### Buffer optimization

In cell-hashing experiments (*10*), staining is performed with a PBS-based staining buffer (SB: 2% BSA, 0.02% Tween-20 in PBS). We initially used this buffer during staining for nucleus hashing as well (gender-specific expression and species-mixing experiments) (*10*). To further optimize our protocol, we compared both a PBS-based staining buffer and a Tris-based staining buffer (ST-SB, see **protocol** below, 2% BSA, 0.02% Tween-20, 10mM Tris, 146mM NaCl, 1mM CaCl_2_, 21mM MgCl_2_) to a non-hashed control observing better performance in ST-SB, in terms of overall agreement with non-hashed controls and in the number of genes recovered per nucleus (**Supplementary Figure 1**). We therefore recommend to perform the staining and washing steps of nucleus-hashing in ST-SB (see detailed **Protocol** below).

### SnRNA-Seq data analysis

Starting from BCL files obtained from Illumina sequencing, we ran cellranger mkfastq to extract sequence reads in FASTQ format, followed by cellranger count to generate gene-count matrices from the FASTQ files. Since our data are from single nuclei, we built and aligned reads to genome references with pre-mRNA annotations, which account for both exons and introns. Pre-mRNA annotations improve the number of detected genes significantly compared to a reference with only exon annotations (*15*). For human and mouse data, we used the GRCh38 and mm10 genome references, respectively. To compare samples of interest (*e.g.*, different loading concentrations), we pooled their gene-count matrices together, and filtered out low-quality nuclei identified based on any one of the following criteria: (**1**) a total number of expressed genes < 200; (**2**) a total number of expressed genes >= 6,000; or (**3**) a percentage of RNA UMIs from mitochondrial genes >= 10%. We performed dimensionality reduction, clustering and visualization on the filtered count matrix as previously described (*16, 17*). Specifically, we selected highly variable genes as previously described (*18*) with a z-score cutoff at 0.5, performed PCA and selected the top 50 principal components (PCs) (*19*), clustered the data based on the 50 selected PCs using the Louvain community detection algorithm (*20*) with a resolution at 1.3. We identified cluster-specific gene expression by differential expression analyses between nuclei within the cluster and outside of the cluster (*16*) using Welch’s t-test and Fisher’s exact test; controlled false discovery rates (FDR) at 5% using the Benjamini-Hochberg procedure (*21*), and annotated putative cell types based on legacy signatures of human and mouse brain cells. We visualized the reduced dimensionality data using tSNE (*22*) with perplexity at 30. Note that in experiments 1 and 4 (**Supplementary Table 1**), we identified one cluster that did not express any known cell type markers and had the lowest median number of RNA UMIs among all clusters. We removed it from further analysis, and repeated the above analysis workflow, except the low-quality nucleus filtration step.

### DemuxEM

Suppose we multiplex *n* samples together. For each droplet, we have a count vector of hashtag UMIs from each sample, (*c_1_*, ⋯, *c_n_*). Each hashtag UMI in the vector can either originate from a properly stained nuclear pore complex (signal) or come from ambient hashtag UMIs (background). We define Θ = (*θ_0_, θ_1_, ⋯, θ_n_*), where θ_0_ is the probability that a hashtag UMI is from the background, and θ_1_, ⋯, θ_n_ are the probabilities that the hashtag UMI is true signal 1,⋯, *n*. If a hashtag UMI is from the background, we denote *P* = (*p*_1_, ⋯, *p*_n_) as the probabilities that this hashtag UMI matches the barcode sequence of samples 1, ⋯, *n*. In addition, we require 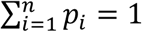.

The probability of generating a hashtag UMI that matches sample *i*’s barcode sequence is:

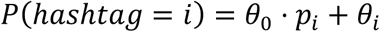

And the log-likelihood of generating the hashtag UMI vector is:

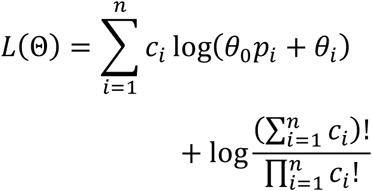

DemuxEM estimates two sets of parameters: (**1**) the background distribution *P* = (*p*_1_, ⋯, *p*_n_), and (**2**) Θ = (θ,, θ_1_, ⋯, θ_n_).

We estimate the background distribution using empty droplets. To identify empty droplets, we first collect all bead barcodes with at least one hashtag UMI. We then calculate the total number of hashtag UMIs each collected bead barcode has and performed a K-means clustering with k = 2 on the total hashtag UMIs. The cluster with a lower mean hashtag UMI number was identified as empty droplets. If we denote the set of identified empty droplets as *B*, we can estimate the background distribution as follows:

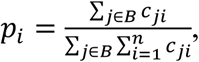

where *c_ji_* is the number of hashtag UMIs matching sample *i* in bead barcode *j*.

We estimate Θ using an Expectation-Maximization algorithm. First, we impose a sparse Dirichlet prior on Θ, Θ~*Dir*(1, 0, ⋯, 0), to encourage the background distribution to explain as much hashtag UMIs as possible. We then follow the EM procedure below:

– E step:

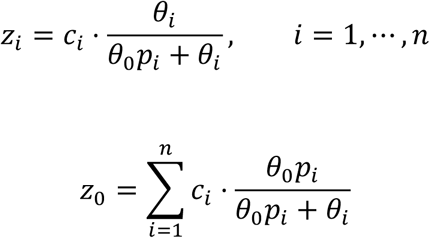

– M step:

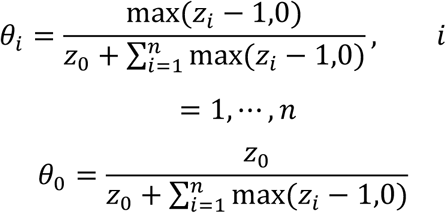

Once we have Θ estimated, we first calculate the expected number of signal hashtag UMIs:

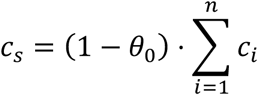

If *c*_s_ < 10, the hashtag UMI vector contains too little signal and thus we mark this droplet as ‘unassigned’. Otherwise, we count the number of samples that has at least 10% signal hashtag UMIs, 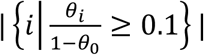. If this number is 1, the droplet is a singlet. Otherwise, it is a multiplet.

### Generation of the “species-mixing” plot

We only used RNA UMIs that were confirmed by at least 2 reads to generate the “species-mixing” plot. Requiring at least 2 reads to confirm a RNA UMI helps filter potentially erroneous RNA UMIs produced from PCR and sequencing errors.

### Estimation of cost per single nucleus in the

We estimate the reduction in cost per single nucleus for a given pricing structure, assuming *X* for loading one 10X channel, *Y* for sequencing one HiSeq lane, and *Z* for the TotalSeq nuclei hashtag cost per hashed sample, to allow readers to determine the costs for their own pricing structures. We sequenced 4 HiSeq lanes in total for four overloading experiments, with proportions roughly as 1: 3: 6: 9 (500 nuc/µl:1,500 nuc/µl:3,000 nuc/µl:4,500 nuc/µl). Based on these values, the sequencing costs for the four settings are 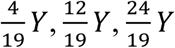 and 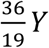 respectively. Adding the 10X channel cost of *X*, and the TotalSeq nuclei hashtag costs of 8*Z*, the final cost for each setting is 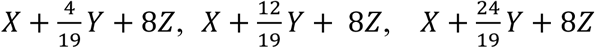 and 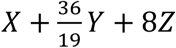 respectively. We then divide each cost by the total number of singlets we detected (**Supplementary Table 3**) to obtain cost per single nucleus in each overloading setting.

## Detailed Protocol

### Materials

| NAME | CATALOG # | VENDOR |
| --- | --- | --- |
| BSA-Molecular Biology Grade - 12 mg | B9000S | New England Biolabs |
| Dounce homogenizers | D8938-1SET | Sigma |
| Pre-Separation Filters (30 µm) | 130-041-407 | Miltenyi Biotec |
| Pre-Separation Filters (20 µm) | 130-101-812 | Miltenyi Biotec |
| Eppendorf® LoBind microcentrifuge tubes | Z666505-100EA | Sigma Aldrich |
| Human TruStain FcX™ | 422302 | BioLegend |
| Beckman Coulter SPRI SELECT REAGENT 5ML | NC0406406 | Fisher Scientific |
| KAPA HiFi HotStart ReadyMix | NC0465187 | Fisher Scientific |

#### 1 Prepare buffers fresh

NP40 Lysis Buffer (NST): 0.1% NP40, 10mM Tris, 146mM NaCl, 1mM CaCl_2_, 21mM MgCl_2_, 40U/mL of RNAse inhibitor

ST Wash Buffer: (10mM Tris, 146mM NaCl, 1mM CaCl_2_, 21mM MgCl_2_), 0.01% BSA (NEB B9000S), 40U/mL of RNAse inhibitor

ST Staining buffer (ST-SB): 2%BSA, 0.02%Tween-20, 10mM Tris, 146mM NaCl, 1mM CaCl_2_, 21mM MgCl_2_)

#### 2 Tissue lysis and homogenizing

Nuclei were extracted as previously described (*1*) with the following minor modifications:

a) For each sample to barcode and pool: prepare a separate homogenizer and douncing pestles A & B. Add 1ml NST buffer to the dounce homogenizer and keep on ice.

Note: Keep tissues/homogenate and buffers on ice throughout the protocol. Pre-cool the centrifuge to 4C and keep at 4C for all steps.

b) Cut a 50-200mg section of frozen brain tissue with a scalpel and dissect to remove white matter and vasculature. Mince tissue and add it to the homogenizer.
c) with a total volume of 1mL, dounce 20 times with pestle A followed by 20 times with pestle B.
d) Add 1ml of ST buffer, filter through 30µm filters (Milentyi Biotec 130-041-407) and transfer filtered homogenate to a 15mL tube.
e) Rinse the homogenizer with 3x 1ml of ST buffer, filter through 30µm filters and add to the filtered homogenate to add up to a final volume of 5ml.
f) Immediately spin down at 500g for 5 mins at 4C to pellet the nuclei in swing bucket rotor
g) Remove supernatant
h) Resuspend nuclei in 200µl of ST-SB, filter with 20um (miltenyibiotec 130-101-812) and transfer to a lo-bind 1.5ml tube (Sigma-Aldrich, Z666505-100EA)

### Count nuclei

Nuclei were counted using the Nexcelom Cellometer Vision 10x objective and a DAPI stain.

a) DAPI was diluted to 2.5µg/µl in ST Buffer.
b) 20µl of the DAPI was pipet mixed with 20ul of the nuclei suspension and 20µl was loaded onto a cellometer cell counting chamber of standard thickness (Nexcelom catalog number: CHT4-SD100-002) and counted using a custom assay with the dilution factor set to 2.

### Hashtag antibody staining

Note: this part mirrors the cell-hashing protocol (*10*), with very minor differences.

a) Add 10 µl Fc Blocking reagent (Biolegend 422302) per 1-2M of nuclei in 100µl of ST-SB/nuclei and incubate for 5 minutes at 4C.
b) Add 1 µg of single nuclei hashing antibody per 100µl of ST-SB/nuclei mix and incubate for 10 minutes at 4C.
c) Wash nuclei 3 times with 1.2 mL ST-SB, spin in swinging bucket rotor for 5 minutes at 500g and 4°C.
d) Resuspend nuclei in ST-SB at 500-3,000 cells/µl.
e) Filter nuclei through MACS Pre-Separation Filters (20µm), and count nuclei to verify concentration after filtration. Adjust to desired concentration.
f) Pool all samples at desired proportions and immediately proceed to next step.

### 10X Genomics single-nuclei sequencing

Use 14µl of pooled sample as input into the 10X Genomics single-cell 3’ v2 assay and process as described until before cDNA amplification.

### Library preparation

a) To increase yield of HTO products during the 10X Genomics cDNA amplification step: Add 1µl of 2µM HTO PCR additive primer (5’GTGACTGGAGTTCAGACGTG TGC*T*C)
b) After cDNA amplification: Separate HTO-derived cDNAs (<180bp) and mRNA-derived cDNAs (>300bp). Perform SPRI selection to separate mRNA-derived and antibody-oligo-derived cDNAs. DO NOT DISCARD SUPERNATANT FROM 0.6X SPRI. THIS CONTAINS THE HASHTAGS.
c) Add 0.6X SPRI (Beckman Coulter, B23317) to cDNA reaction as described in 10X Genomics protocol.
d) Incubate 5 minutes and place on magnet. Supernatant contains hashtags, and beads contain full length mRNA-derived cDNAs.

### Library preparation for mRNA-derived cDNA >300bp (bead fraction)

Proceed with standard 10X protocol for cDNA sequencing library preparation.

### Library preparation for mRNA-derived cDNA <300bp (supernatant fraction)

Purify Hashtags using two 2X SPRI purifications per manufacturer protocol:

- Add 1.4X SPRI to supernatant to obtain a final SPRI volume of 2X SPRI.
- Transfer entire volume into a low-bind 1.5mL tube.
- Incubate 10 minutes at room temperature.
- Place tube on magnet and wait ~2 minutes until solution is clear.
- Carefully remove and discard the supernatant.
- Add 400 µl 80% ethanol to the tube without disturbing the pellet and stand for 30 seconds (only one ethanol wash).
- Carefully remove and discard the ethanol wash.
- Centrifuge tube briefly and return it to magnet.
- Remove and discard any remaining ethanol.
- Resuspend beads in 50 µl water.
- Perform another round of 2X SPRI purification by adding 100 µl SPRI reagent directly onto resuspended beads.
- Mix by pipetting, and incubate 10 minutes at room temperature.
- Place tube on magnet and wait ~2 minutes until solution is clear.
- Carefully remove and discard the supernatant.
- Add 200 µl 80% ethanol to the tube without disturbing the pellet and let stand for 30 seconds (first Ethanol wash).
- Carefully remove and discard the ethanol wash.
- Add 200 µl 80% ethanol to the tube without disturbing the pellet and let stand for 30 seconds (second Ethanol wash).
- Carefully remove and discard the ethanol wash.
- Centrifuge tube briefly and return it to magnet.
- Remove and discard any remaining ethanol and allow the beads to air dry for 2 minutes (do not over-dry beads).
- Resuspend beads in 90 µl water.
- Mix vigorously by pipetting and incubate at room temperature for 5 minutes.
- Place tube on magnet and transfer clear supernatant into PCR well.
- Prepare 100µL PCR reaction with purified small fraction:

o45 µl purified Hashtag fraction

o50 µl 2x KAPA Hifi PCR Master Mix.

o2.5 µl TruSeq DNA D7xx_s primer (containing i7 index) 10 µM. (i.e. D701: 5’CAAGCAGAAGACGGCA

TACGAGATCGAGTAAT GTGACTGGAGTTCAGACGTGT* G*C)

o2.5 µl SI PCR oligo at 10 µM (SI PCR: 5’AATGATACGGCGACCACCGAGATCTACACTCTTTCCCT ACACGACGC*T*C)

Cycling conditions: 95°C 3 min

95°C 20 sec |

64°C 30 sec | ~ 8 cycles

72°C 20 sec |

72°C 5 min

Perform 1.6X SPRI purification by adding 160 µl SPRI reagent.

- Incubate 5 minutes at room temperature.
- Place tube on magnet and wait 1 minute until solution is clear.
- Carefully remove and discard the supernatant.
- Add 200 µl 80% ethanol to the tube without disturbing the pellet and let stand for 30 seconds (first ethanol wash).
- Carefully remove and discard the ethanol wash.
- Add 200 µl 80% ethanol to the tube without disturbing the pellet and let stand for 30 seconds (second ethanol wash).
- Carefully remove and discard the ethanol wash.
- Centrifuge tube briefly and return it to magnet.
- Remove and discard any remaining ethanol and allow the beads to air dry for 2 minutes.
- Resuspend beads in 20 µl water.
- Pipette mix vigorously and incubate at room temperature for 5 minutes.
- Place tube on magnet and transfer clear supernatant to PCR tube.

### Quantify library

Quantify library by standard methods (QuBit, BioAnalyzer). Hashtag library will be around 180 bp.

### Sequence

Combine mRNA library and HTO library (~90% mRNA to 10% HTO), and sequence with the regular 10X RNA-seq read structure:

oRead 1 = 26

oRead 2 = 55 bp

oIndex 1 = 8 bp

oIndex 2 = n/a

---

**Figure S1.**
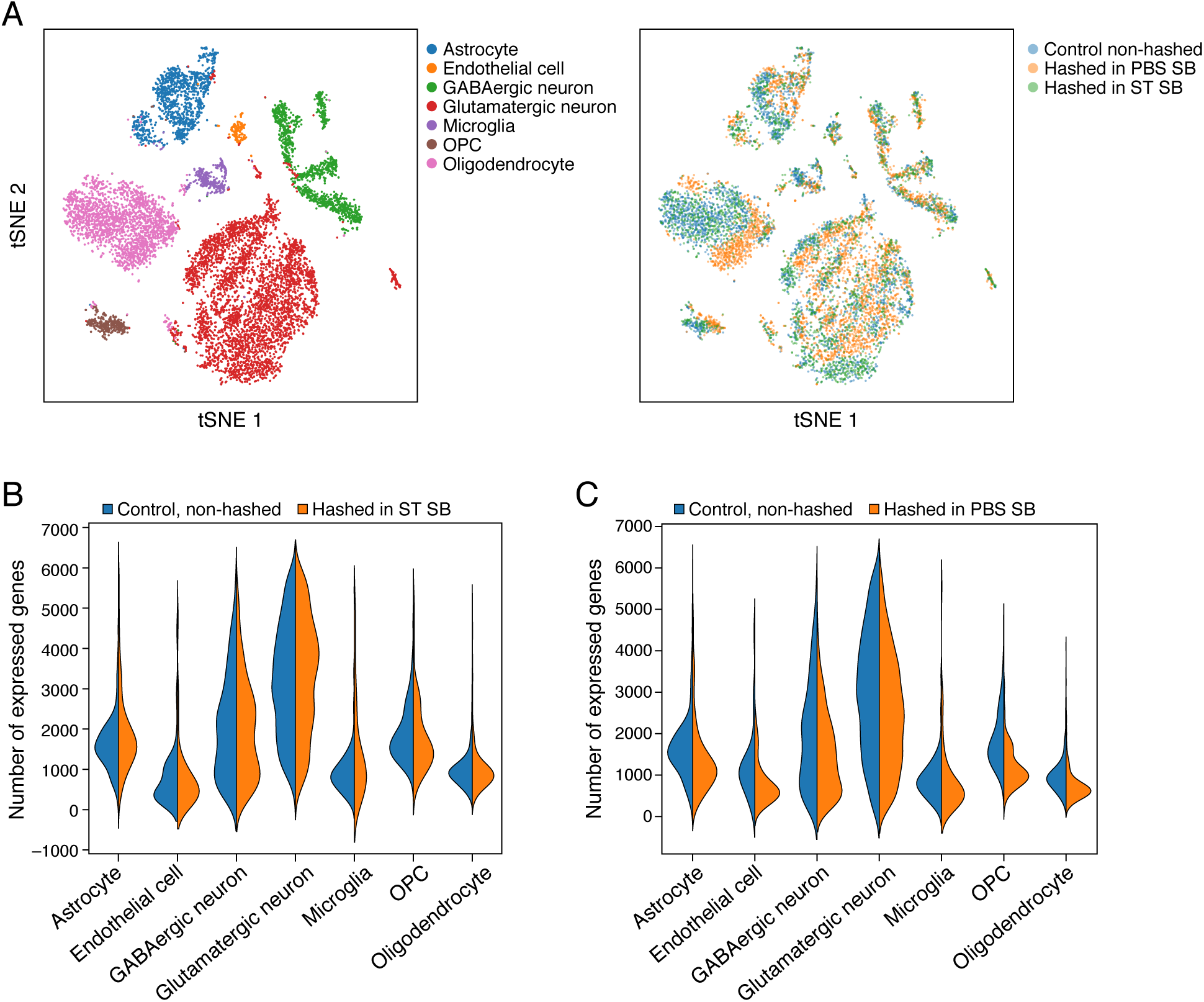
Buffer optimization for multiplexing. **a.** tSNE of single nucleus profiles from non-hashed control, PBS-based (PBS-SB) and ST-based staining buffer (ST-SB) colored by either cell type (left) or protocol (right). Nuclei stained with ST-SB buffer (green) largely overlap with the non-hashing control nuclei (blue), whereas PBS-stained nuclei (orange) show some separation within the clusters. **b,c.** Decreased number of expressed genes detected when using PBS-SB. Distribution of number of expressed genes (*y* axis) across cell types (*x* axis) for nuclei stained with ST-SB (**b**, orange) or PBS-SB (**c**, orange) compared to the non-hashing control (blue). Except for microglia, ST-SB performs better across cell types.

**Figure S2.**
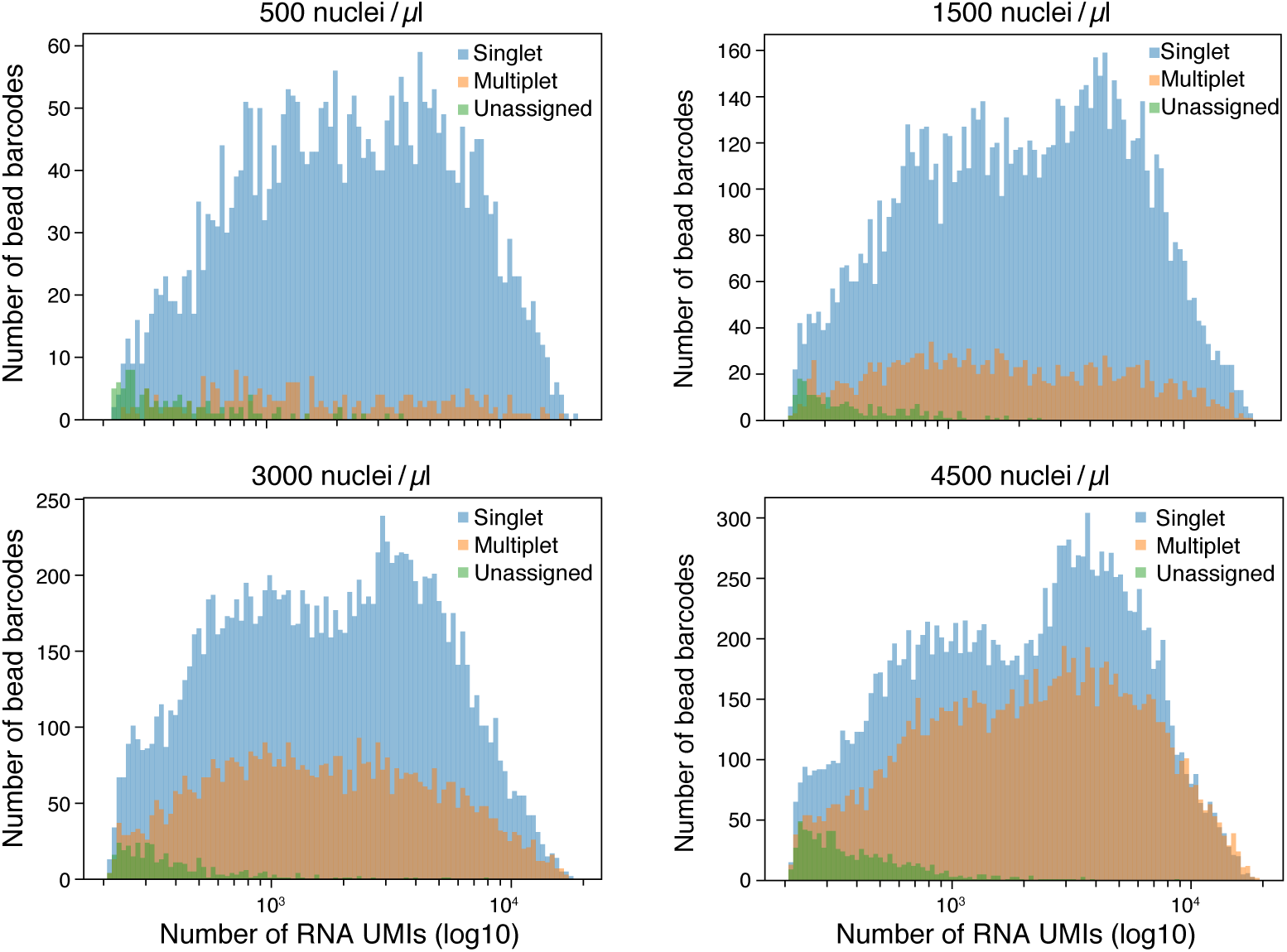
Nuclei multiplets do not necessarily have a larger number of RNA UMIs. Distribution of number of bead barcodes (*y* axis) for beads with different numbers of detected UMIs (*x* axis), for singlets (blue), multiplets (orange) and unassigned droplets (green), in 8 hashed human cortex samples loaded at concentrations of 500, 1,500, 3,000 or 4,500 nuclei/μl. Although the multiplet rate rises with increasing loading concentrations, we observe similar RNA UMI count distributions for multiplets and singlets, a feature not observed for single-cell hashing (*10*).

